# Whole-genome sequences suggest long term declines of spotted owl (*Strix occidentalis*) (Aves: Strigiformes: Strigidae) populations in California

**DOI:** 10.1101/343434

**Authors:** Zachary R. Hanna, John P. Dumbacher, Rauri C.K. Bowie, Jeffrey D. Wall

## Abstract

We analyzed whole-genome data of four spotted owls (*Strix occidentalis*) to provide a broad-scale assessment of the genome-wide nucleotide diversity across *S. occidentalis* populations in California. We assumed that each of the four samples was representative of its population and we estimated effective population sizes through time for each corresponding population. Our estimates provided evidence of long-term population declines in all California *S. occidentalis* populations. We found no evidence of genetic differentiation between northern spotted owl (*S. o. caurina*) populations in the counties of Marin and Humboldt in California. We estimated greater differentiation between populations at the northern and southern extremes of the range of the California spotted owl (*S. o. occidentalis*) than between populations of *S. o. occidentalis* and *S. o. caurina* in northern California. The San Diego County *S. o. occidentalis* population was substantially diverged from the other three *S. occidentalis* populations. These whole-genome data support a pattern of isolation-by-distance across spotted owl populations in California, rather than elevated differentiation between currently recognized subspecies.

## Introduction

Spotted owls (*Strix occidentalis*) in California are an important protected species that is declining throughout most of its range. Their status has enormous economic implications (Montgomery *et al.* 1994). The species is represented in California by two recognized subspecies, the northern spotted owl (*S. o. caurina*) and the California spotted owl (*S. o. occidentalis*). In California, *S. o. caurina* occurs from the northern border with Oregon, south along the west side of California’s Central Valley to the Golden Gate strait, and southeast to the region between the Pit River and Lassen Peak. The latter region constitutes the dividing line between *S. o. caurina* and *S. o. occidentalis* (Barrowclough *et al.* 2011; Tempel *et al.* 2017). *Strix occidentalis occidentalis* occurs from this line south to the southern Sierra Nevada and in mountainous areas of southern California, including the coastal ranges west of the Central Valley as far north as the Santa Lucia Mountains and Monterey Bay (Barrowclough *et al.* 1999, 2011 p. 2017; Davis and Gould 2008; Gutiérrez *et al.* 2017). *Strix occidentalis caurina* currently has a threatened listing status under the United States of America Endangered Species Act as well as the California Endangered Species Act. *Strix occidentalis occidentalis* is in decline throughout much of its range (Tempel *et al.* 2017). Although not presently listed under either endangered species act, *S. o. occidentalis* is currently designated by the California Department of Fish and Wildlife as a Species of Special Concern (Davis and Gould 2008).

There have been multiple excellent studies of the population genetics of *Strix occidentalis caurina* and *S. o. occidentalis*. Most of these relied on mitochondrial DNA (Barrowclough *et al.* 1999, 2005, 2011; Haig *et al.* 2004), which comprises a single genetic locus that does not always accurately represent the history of the entire nuclear genome. Additional studies analyzed microsatellites (Funk *et al.* 2008), RAPD markers (Haig *et al.* 2001), or a combination of mitochondrial DNA and microsatellites (Miller *et al.* 2017), but these mostly focused on *S. o. caurina* and northern populations of *S. o. occidentalis*. A whole genome analysis would test whether earlier conclusions based on mitochondrial genes and several nuclear markers are generalizable. By sequencing whole genomes, we can acquire data for thousands of independent loci and, even with one individual per population, we can obtain information for thousands of additional samples of the evolutionary history of a population as compared with previous *S. occidentalis* genetic studies.

The barred owl (*Strix varia*) has invaded the entire range of *Strix occidentalis caurina* and is in contact with *S. o. occidentalis* from the Lassen area to the southern Sierra Nevada (Keane 2017). *Strix occidentalis* and *S. varia* hybridize, but the frequency and primary directionality of introgression is unknown. In order to detect introgression on the leading edge of the *S. varia* expansion in California, we need a better understanding of the genetic variation present in pure *S. occidentalis* populations.

The recently assembled genome of a *Strix occidentalis caurina* individual from Marin County, California (Hanna *et al.* 2017d, 2017c) provides a useful resource for characterizing the genomic variation in *S. occidentalis* throughout California. Here we report data from high-coverage whole-genome sequences of that individual and new sequences of three additional *S. occidentalis*. Our sampling spans the northern and southern geographic extremes of *S. o. caurina* and *S. o. occidentalis* within California (Figure 1). We use these data to provide a broad-scale assessment of the genomic divergence of *S. occidentalis* in California and estimate long-term trends in population size.

**Figure 1.**
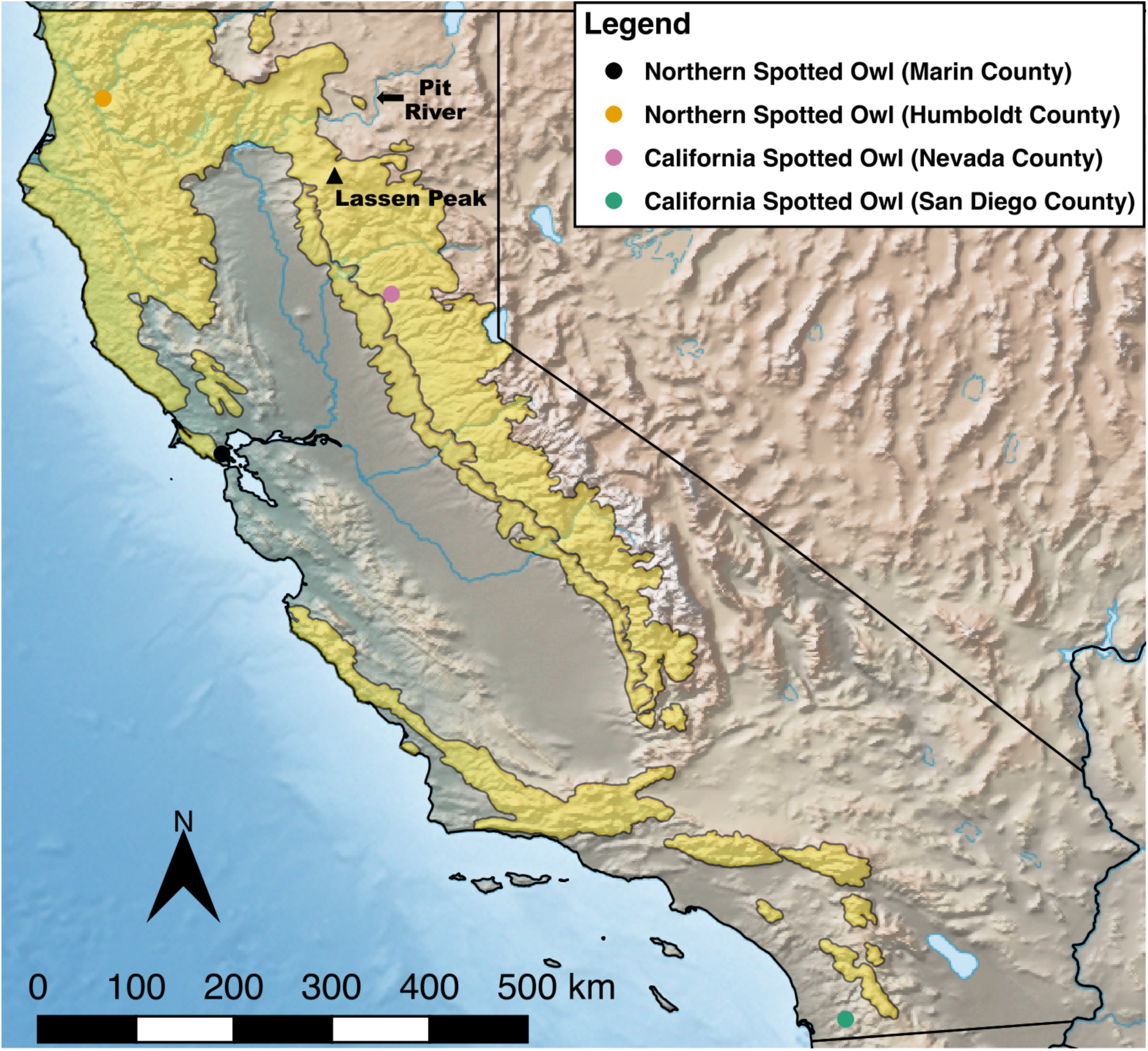
California map of the collection locations of the *Strix occidentalis* specimens from which we obtained whole-genome sequences for this study. The yellow shading illustrates the distribution of *Strix occidentalis* in California. We have indicated the locations of the Pitt River and Lassen Peak. Owls to the north of the region between these two geographic features are considered northern spotted owls (*S. o. caurina*) while those to the south of this region are California spotted owls (*S. o. occidentalis*) (Barrowclough *et al.* 2011; Tempel *et al.* 2017).

## Results

The nuclear reference genome contained 1,113,365,877 non-N nucleotides after masking. Our final set of variants included 486,582 biallelic sites. Across individuals, the mean coverage at the final set of variant sites ranged from approximately 33.2 to 56.7X (Table S1). Grouping the results by subspecies, we recovered a higher nucleotide diversity within *Strix occidentalis occidentalis* (1.676 × 10^−4^) than *S. o. caurina* (1.454 × 10^−4^) in California (Table 1). We calculated an *F_ST_* of 0.1727 for the two subspecies. When we treated each sample as a separate population, additional patterns emerged.

We found no divergence (*F_ST_* = 0) between northern spotted owl populations in California. The Marin County population had a lower nucleotide diversity (1.374 × 10^−4^) than the Humboldt County population (1.591 × 10^−4^); between them, these two populations had the lowest nucleotide diversity (1.399 × 10^−4^) of any population pair (Table 1).

**Table 1.**
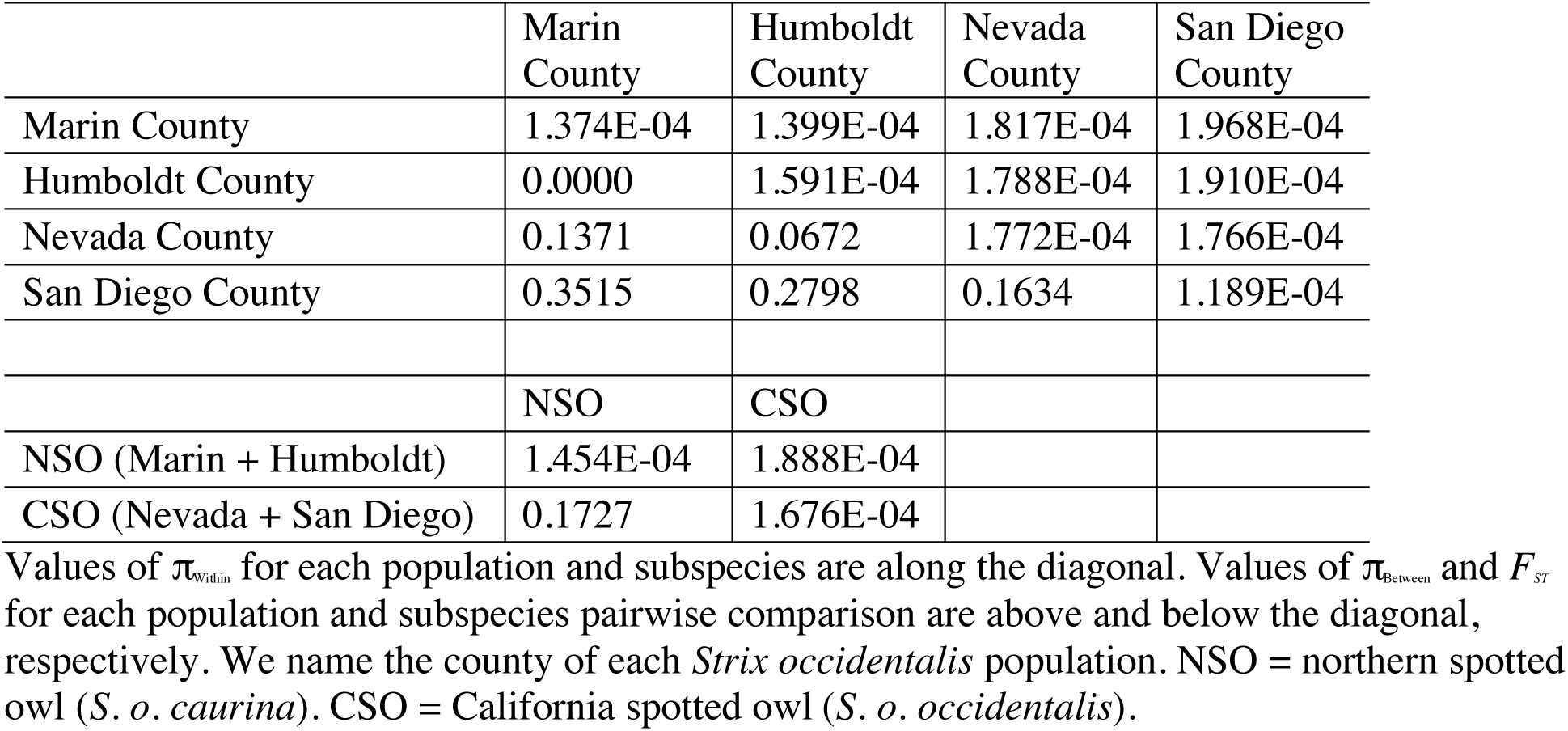
Population and subspecies diversity and divergence statistics.

The San Diego County population harbored the lowest nucleotide diversity of all of the populations (1.189 × 10^−4^). We calculated genome-wide *F_ST_* values ranging from 0.1634 to 0.3515 for pairwise comparisons of the San Diego County population with the other three populations (Table 1). The lowest of these (0.1634) was the *F_ST_* between San Diego County and Nevada County. However, we found lower differentiation between Nevada County and each of the *Strix occidentalis caurina* populations (*F_ST_* = 0.0672 between Nevada and Humboldt County; *F_ST_* = 0.1371 between Nevada and Marin County) than between the Nevada and San Diego County populations of *S. o. occidentalis*.

Across all individuals, the proportion of the genome comprised by runs of homozygosity (ROH) ranged from approximately 6.08-36.23%, with the lowest proportion found in the Humboldt County population and the highest in the San Diego County population (Table S2). We found the two longest tracts in the Marin and San Diego County populations (7.5 and 8.9 Mb, respectively) (Figure 2). As the majority of the tracts across all individuals were less than 4 Mb long and none of the ROH exceeded 10 Mb (Figure 2), we found no evidence for very recent inbreeding (e.g. within the last three generations). Rather, the frequency and length of the ROH suggest low historic population sizes, especially in the Marin and San Diego County populations.

**Figure 2.**
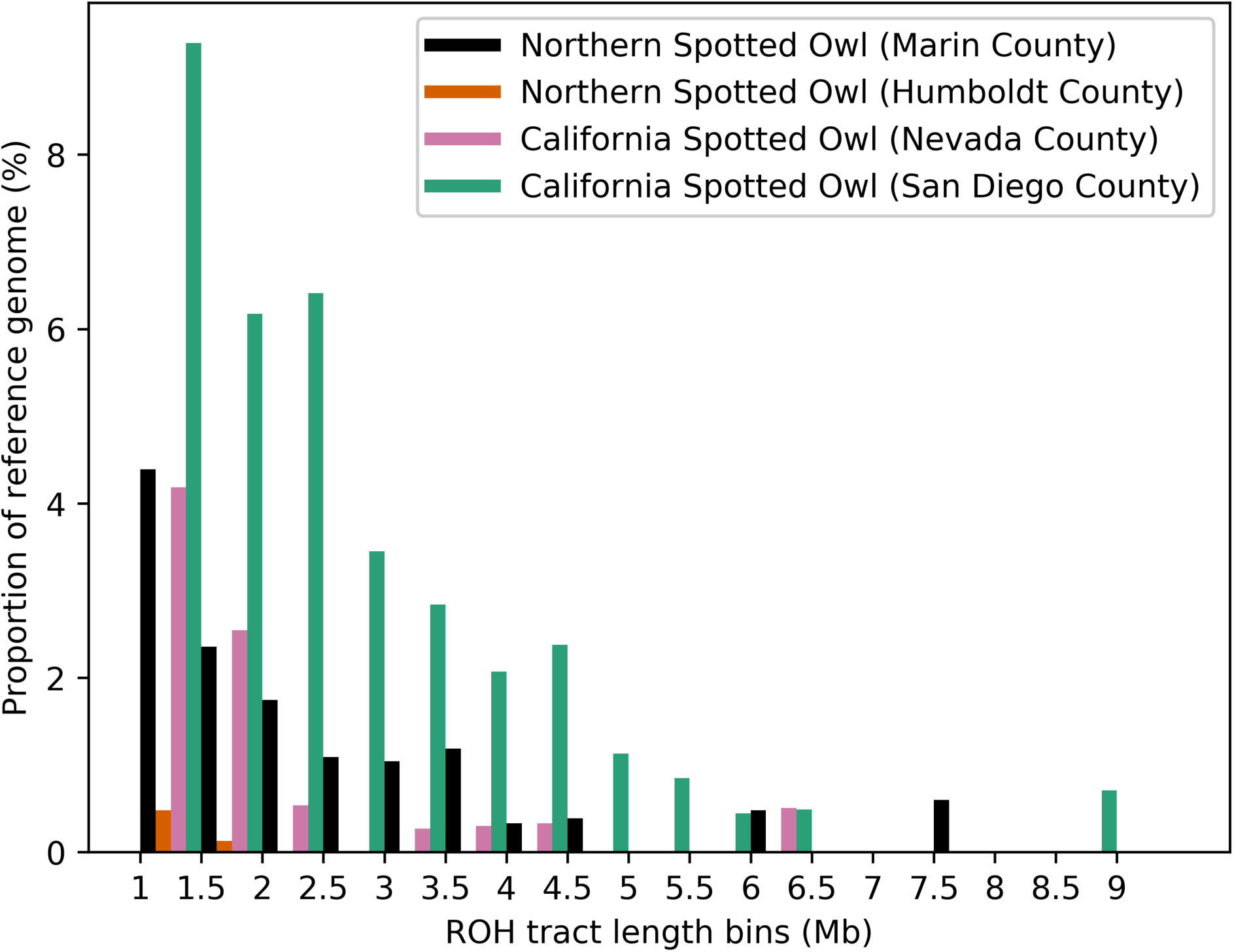
Histogram of runs of homozygosity. The x-axis provides length bins for runs of homozygosity (ROH) in each population. Each bin contains ROH greater than or equal to the length in megabases (Mb) indicated by the left tick mark and less than the length of the right tick mark. The y-axis expresses the total length of all ROH in a length bin as a percentage of the length of the reference genome.

Our estimates of the effective population size (*N_e_*) of each population indicate that all four populations have undergone long-term population declines over the past approximately 70,000 generations. Of note, the *N_e_* of the *Strix occidentalis caurina* populations grew about an order of magnitude larger approximately 3,000 to 700 generations ago before declining again (Figure 3). Taken with the caveat that recent population size changes are difficult to estimate accurately using the SMC++ method or other similar methods, especially with low sample sizes (Terhorst *et al.* 2017), our most recent estimates suggest that the San Diego County population has the lowest current *N_e_* (~ 350 individuals) and that of the Marin County population (~ 500 individuals) is only slightly higher. We estimated the current *N_e_* of the Nevada County population at ~ 1,100 individuals. The Humboldt County population *N_e_* (~ 4,900 individuals) is approximately one order of magnitude greater than that of both the Marin and San Diego County populations. The *N_e_* of the *S. o. caurina* populations appear to have remained very similar until the last 700 to 800 generations when they diverged; the Marin County effective population size dropped by over an order of magnitude while the Humboldt County population declined less drastically. The *N_e_* of the *S. o. occidentalis* populations has also been similar over time, but less so than that of the *S. o. caurina* populations. The San Diego County population has been undergoing a steady decline in *N_e_* for approximately the past 8,000 generations.

**Figure 3.**
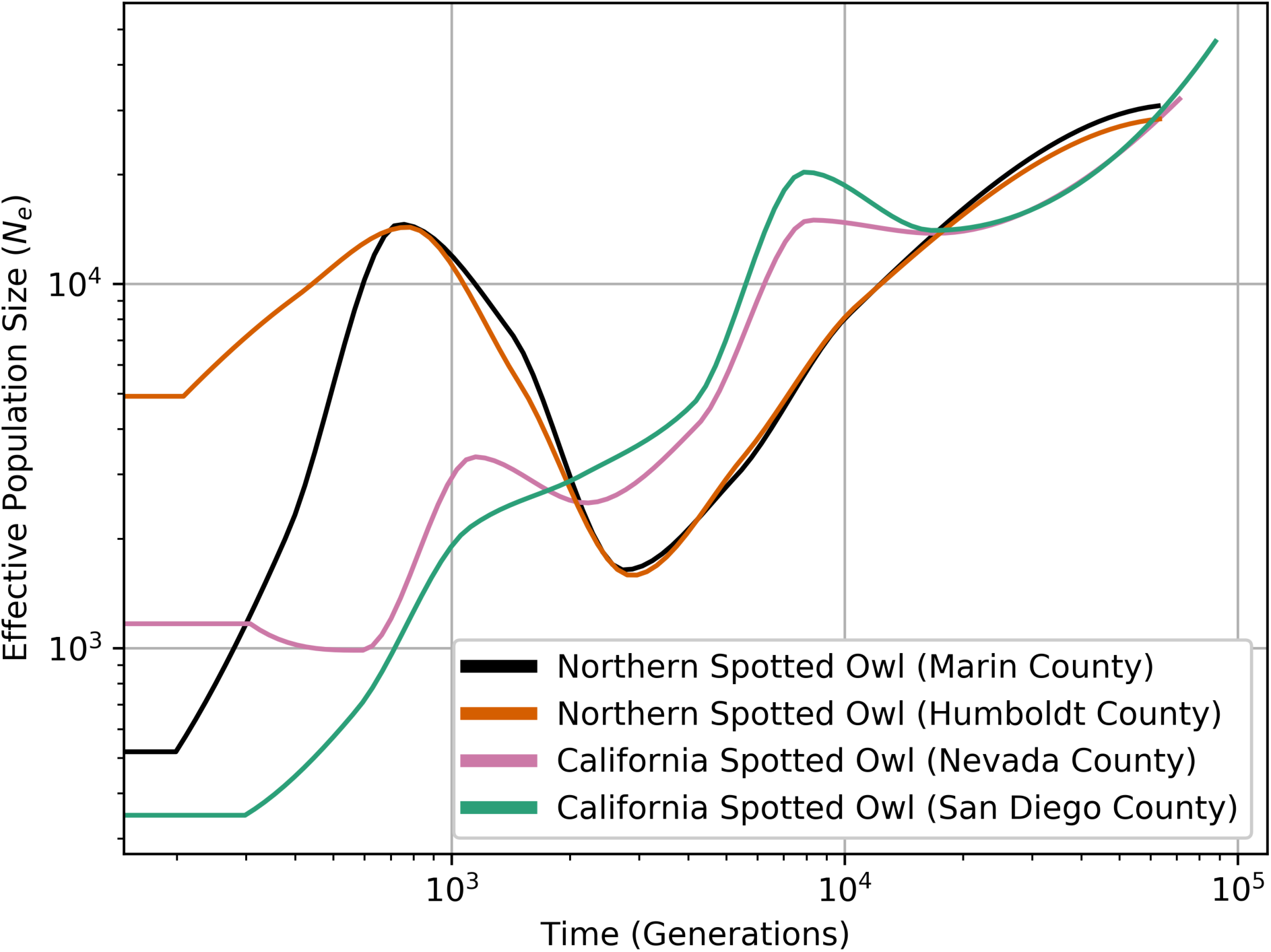
Effective population size through time of each *Strix occidentalis* population. The time on the x-axis is in generations before present. Both the x and y axes are logarithmically scaled.

## Discussion

Early genetic studies of spotted owls (*Strix occidentalis*) utilizing mitochondrial DNA (mtDNA) sequence data concluded that *S. o. caurina* and *S. o. occidentalis* are largely reciprocally monophyletic, but that there are migrants across the subspecies’ boundaries and evidence of incomplete lineage sorting (Barrowclough *et al.* 1999, 2005; Haig *et al.* 2004). Funk et al. (2008) utilized a panel of ten nuclear-genome-derived microsatellite loci and they found two genetic clusters nearly exclusive to *S. o. caurina* and a third in which all of their *S. o. occidentalis* samples had high membership, but their results also included many *S. o. caurina* with membership in this third cluster. They interpreted their results as supporting the current division of subspecies, but with substantial gene flow between *S. o. caurina* and *S. o. occidentalis* in northern California and southern Oregon (Funk *et al.* 2008). Studies with denser sampling in northern California near the *S. o. caurina* - *S. o. occidentalis* boundary found multiple *S. o. caurina* genetic clades, which they attributed to incomplete lineage sorting, along with evidence of a differentiated *S. o. caurina* population in northern California (Barrowclough *et al.* 2011; Miller *et al.* 2017). The Miller et al. (2017) investigators further concluded that gene flow is predominantly occurring from the range of the *S. o. occidentalis* into that of *S. o. caurina*.

We found higher nucleotide diversity in the Nevada County *Strix occidentalis occidentalis* population than in the Humboldt County *S. o. caurina* population. This contrasts with the findings of prior researchers, who have reported higher nucleotide diversity in Humboldt County *S. o. caurina* than in Sierra Nevada *S. o. occidentalis* (Barrowclough *et al.* 1999; Miller *et al.* 2017), although Barrowclough et al. (2005) found higher diversity in Lassen County than in both Humboldt County *S. o. caurina* and El Dorado County *S. o. occidentalis*. El Dorado County is approximately 90 km southeast of the location of our Nevada County sample. Higher genetic diversity in Nevada County was contrary to what we expected if gene flow was predominantly northward out of the range of *S. o. occidentalis* and into that of *S. o. caurina* in northern California, as Miller et al. (2017) suggested. Indeed, the nucleotide diversity of the Nevada County population was the highest of all four of our populations.

In contrast with our finding of the highest genetic diversity in the Nevada County population, we estimated that the effective size of the Nevada County population has been lower than that of the Humboldt County population for approximately the past two thousand generations. The Marin County and San Diego County populations, each of which represents the southern edge of the range of their respective subspecies [although an isolated *Strix occidentalis occidentalis* population exists south of San Diego County in the Sierra de San Pedro Mártir of Baja California (Bryant 1889)], have low genetic diversity and low recent effective population sizes. All population sizes are lower than their historic counterparts. We estimated that the Humboldt County population size has declined the least in comparison with historic levels. Both *S. o. occidentalis* populations appear to have been declining steadily with little to no evidence of periodic recovery. In contrast, the *S. o. caurina* populations show indication of a period of sustained population growth from approximately 3,000 to 700 generations ago. Depending on whether one uses two (Gutiérrez *et al.* 1995), five (Barrowclough and Coats 1985; Barrowclough *et al.* 1999), or ten years (Noon and Biles 1990; USDA Forest Service 1992) as the *S. occidentalis* generation time, the *S. o. caurina* effective population size trough and subsequent growth 3,000 generations ago could coincide roughly with the beginning of the retreat of glaciers in North America approximately 19,000-20,000 years ago after the last glacial maximum (Clark *et al.* 2009). However, it is important to consider that recent population crashes in some or all of these populations (Gutiérrez 1994; Davis and Gould 2008; Dugger *et al.* 2015; Tempel *et al.* 2017) may be affecting our ability to estimate historic population sizes accurately (Terhorst *et al.* 2017).

Our pairwise *F_ST_* value between the populations grouped by subspecies appeared to be driven largely by the divergence of the Marin and Humboldt County *Strix occidentalis caurina* populations from the San Diego County *S. o. occidentalis* population. We estimated a higher genetic divergence between the Nevada and San Diego County *S. o. occidentalis* populations than between the Nevada County *S. o. occidentalis* population and each of the *S. o. caurina* populations. Our pairwise *F_ST_* value comparing Nevada and Humboldt County (*F_ST_* = 0.0672) was almost identical to that calculated by Funk et al. (2008) between their *S. o. occidentalis* and Humboldt + Del Norte County populations using 10 microsatellite loci (*F_ST_* = 0.066). Notably, the Funk et al. (2008) *S. o. occidentalis* population grouped samples from the northern and southern end of the Sierra Nevada *S. o. occidentalis* range. Our recovery of a comparable *F_ST_* value with our samples from Nevada and Humboldt counties suggests low differentiation across Sierra Nevada *S. o. occidentalis*, a preliminary finding that needs to be tested with further genomic sequencing.

Our whole-genome data show the San Diego County population being clearly diverged from the other three populations. The genetic diversity of the San Diego population was the lowest of the four. Previous investigators have reported low to no genetic diversity in the mitochondrial control region of *Strix occidentalis occidentalis* populations in southern California (Barrowclough *et al.* 1999, 2005; Haig *et al.* 2004). Haig et al. (2004) found no differentiation (*F_ST_* = 0.00) between *S. o. occidentalis* from the Sierra Nevada and southern California, whereas we found substantial differentiation between *S. o. occidentalis* from Nevada and San Diego counties (*F_ST_* = 0.1634). Our *F_ST_* values for all pairwise comparisons with the San Diego County population exceeded all of the *F_ST_* values calculated by the Funk et al. (2008) microsatellite study investigating the relationships between three spotted owl subspecies, even those involving comparisons of the Mexican spotted owl (*<u>Strix o. lucida</u>*) with *S. o. occidentalis* or *S. o. caurina*. The highest pairwise *F_ST_* value calculated by the Funk et al. (2008) authors (*F_ST_* = 0.141) was between their *S. o. occidentalis* and *S. o. lucida* populations; this was lower than the pairwise value we recovered between our *S. o. occidentalis* populations from Nevada and San Diego counties (*F_ST_* = 0.1634). Our results suggest that the southern California *S. o. occidentalis* population is considerably differentiated from the Sierra *S. o. occidentalis* population.

In contrast with our finding substantial differentiation between the *Strix occidentalis occidentalis* populations, we found no genetic differentiation between the Marin and Humboldt County *S. o. caurina* populations. Studies utilizing mtDNA and microsatellites to compare the genetic divergence of *S. o. caurina* populations in Marin and Mendocino County (the latter located immediately south of Humboldt County in California) reported *F_ST_* values of 0.14327 (Chi 2006) and 0.090 (Henke 2005) for mtDNA and microsatellite markers, respectively. A mtDNA study by Barrowclough et al. (2005) also found the Marin County *S. o. caurina* population to be divergent from other *S. o. caurina* populations. According to our estimate, the effective population size of the Marin County population was almost identical to that of the Humboldt County population until approximately 700 generations ago when the Marin County population appears to have undergone a severe reduction, suggesting that Marin County *S. o. caurina* were part of a much larger interbreeding group until fairly recently.

We purposefully chose samples from the northern extent of the California ranges of *Strix occidentalis caurina* and *S. o. occidentalis*, but outside of previously described *S. o. caurina* and *S. o. occidentalis* contact zones (Barrowclough *et al.* 2011; Miller *et al.* 2017). Plausible explanations for the finding of less divergence between the *S. o. caurina* populations and the Nevada County *S. o. occidentalis* population than between the Nevada and San Diego County *S. o. occidentalis* populations include the following: 1) the Barrowclough et al. (2011) *S. o. caurina* and *S. o. occidentalis* contact zone has shifted southward in the Sierra Nevada and our Nevada County sample was an admixed individual; 2) our Humboldt County sample was the result of admixture of *S. o. caurina* and *S. o. occidentalis* in northern California; 3) alleles are moving beyond the Barrowclough et al. (2011) and Miller et al. (2017) contact zones in northern California into the ranges of *S. o. caurina* and/or *S. o. occidentalis*; and/or 4) whole-genome sequences are picking up variation that was missed by the microsatellites and not present in the mitochondrial genome. We attempted to avoid sampling from the zone of admixture of *S. o. caurina* and *S. o. occidentalis* described by Miller et al. (2017) by sampling approximately 50 km west of the described contact region. Nevertheless, it is possible that our Humboldt County sample was an *S. o. occidentalis* that migrated into the range of *S. o. caurina* or a product of admixture of the two subspecies. Likewise, we attempted to avoid the contact zone described by Barrowclough et al. (2011) by sampling an individual collected approximately 150 km southeast of Lassen Peak, but it is possible that our Nevada County sample was a *S. o. caurina* that migrated into the range of *S. o. occidentalis* or it is the result of admixture of two differentiated subspecies.

If our Humboldt County sample was a *Strix occidentalis occidentalis* that migrated into the *S. o. caurina* range or an introgressed individual, it is difficult to explain why we recovered no genetic differentiation between the Humboldt and Marin County populations, especially given the differentiation of Nevada and Marin counties. Our Marin County sample was from a location approximately 250 km south of the Miller et al. (2017) contact zone. If our Nevada County sample was a *S. o. caurina* that migrated into the range of *S. o. occidentalis*, we would have expected to identify a lower level of genetic differentiation between the Marin and Nevada County samples than that which we detected, given the absence of differentiation of the Marin and Humboldt County *S. o. caurina* populations. If admixture between two previously diverged subspecies is occurring as far south into the range of *S. o. occidentalis* as Nevada County, it is hard to image how the entire 640 km length of the Sierra Nevada (Schoenherr 1992) would not be colonized by admixed individuals within a few generations, since juvenile dispersal distance can exceed 100 km in *S. o. caurina* (Forsman *et al.* 2002). Apart from the lack of divergence between the *S. o. caurina* populations, all other population comparisons with our whole-genome dataset seem to support a pattern of isolation by distance, rather than a distinct boundary between *S. o. caurina* and *S. o. occidentalis*.

## Materials and Methods

### Sequence data

We utilized whole genome sequencing data generated by a previous study (Hanna *et al.* 2017c) from a *Strix occidentalis caurina* sample from Marin County, California (NCBI Sequence Read Archive (SRA) run accessions SRR4011595, SRR4011596, SRR4011597, SRR4011614, SRR4011615, SRR4011616, and SRR4011617 for *S. o. caurina* sample CAS:ORN:98821; Table 2; Tables S3 and S4). We also obtained high-coverage whole genome sequences from another *S. o. caurina* (CAS:ORN:99418) and two *S. o. occidentalis* (samples CDFWN13249 and CDFWN14089) by sending genomic DNA to MedGenome (Foster City, California, U.S.A.) for preparation of whole genome libraries with a single, 8 bp index and a 420 bp insert size. MedGenome sequenced each of these libraries with 150 bp paired-end reads on a HiSeq X Ten (Illumina, San Diego, California, U.S.A.). The raw sequences are available from the NCBI SRA run accessions SRR6048828, SRR6048829, SRR6048848 (Table S4).

**Table 2.** Sample information.

| **Specimen Identifier** | **Genus** | **species** | **subspecies** | **Latitude** | **Longitude** | **County** | **State** | **Collection Date** |
| --- | --- | --- | --- | --- | --- | --- | --- | --- |
| CAS:ORN:98821 | *Strix* | *occidentalis* | *caurina* | 37.929565 | -122.5402 | Marin | California | 26-Jun-2005 |
| CAS:ORN:99418 | *Strix* | *occidentalis* | *caurina* | 41.19592798 | -123.6310486 | Humboldt | California | 8-Aug-2011 |
| CAFWN14089 | *Strix* | *occidentalis* | *occidentalis* | 39.396524 | -120.983486 | Nevada | California | 17-Feb-2014 |
| CDFWN13249 | *Strix* | *occidentalis* | *occidentalis* | 32.733285 | -116.812313 | San Diego | California | 10-Nov-2013 |

We list here the taxonomic identification of each sample, the county and state of origin, and the sample collection date.

Our four *Strix occidentalis* samples were from locations near the extremes of the California distributions of *S. o. caurina*) and *S. o. occidentalis*) (Figure 1): Marin County, the southernmost extent of *S. o. caurina* (sample CAS:ORN:98821); Humboldt County, near the northern extent of *S. o. caurina* (CAS:ORN:99418); San Diego County, the southern extent of *S. o. occidentalis* (CDFWN13249); and Nevada County, near the northern extent of *S. o. occidentalis* (CAFWN14089). Our Humboldt County sample was from a location approximately 50 km northwest of the California and northern spotted owl contact zone identified by Miller et al. (2017).

### Alignment and filtering

We used Trimmomatic version 0.36 (Bolger *et al.* 2014) along with relevant adapter files (Bolger *et al.* 2014; Hanna 2018a) to remove adapter sequences and low quality leading and trailing bases. For all four samples we used BWA-MEM version 0.7.17-r1188 (Li 2013) to align the processed sequences to the *S. o. caurina* nuclear genome “StrOccCau_1.0_nuc.fa” (Hanna *et al.* 2017d, 2017c) with the mitochondrial genome sequence from Hanna et al. (2017a) added to the nuclear genome reference sequence. We merged and sorted the alignments and then marked duplicate sequences using Picard version 2.17.6 (http://broadinstitute.github.io/picard; Accessed 2018 Mar 22).

### SNP calling and filtering

We called variants using the Genome Analysis Toolkit (GATK) version 3.8.0 HaplotypeCaller tool (McKenna *et al.* 2010; DePristo *et al.* 2011; Van der Auwera *et al.* 2013) to produce a genomic variant call format (gVCF) file for each sample. We then used the GATK version 3.8.0 GenotypeGVCFs tool to produce a VCF file containing the variant information for all samples. We used the GATK version 3.8.0 SelectVariants and VariantFiltration tools to filter the variants and remove those of suspect quality. We then followed an adapted version of the GATK guidelines for base quality score recalibration BQSR (https://gatkforums.broadinstitute.org/gatk/discussion/2801/howto-recalibrate-base-qualityscores-run-bqsr; Accessed 2018 Mar 15) and used the GATK version 3.8.0 BaseRecalibrator and PrintReads tools to recalibrate all of the alignment files based on our set of high-quality variants. We performed a new round of variant calling using the GATK version 3.8.0 HaplotypeCaller and GenotypeGVCFs tools. We filtered the variant file to retain only high-quality, biallelic single nucleotide polymorphisms (SNPs) that were not on the mitochondrial genome using the GATK version 3.8.0 SelectVariants and VariantFiltration tools as well as GNU Awk (GAWK) version 4.2.0 (Free Software Foundation 2017).

In order to utilize up-to-date repetitive sequence libraries to identify and remove variants in low-complexity and repetitive regions, we performed a new masking of repetitive sequences in the reference *Strix occidentalis caurina* genome StrOccCau_1.0_nuc.fa (Hanna *et al.* 2017d, 2017c) with the mitochondrial genome from Hanna et al. (2017a). We created a *de novo* model of the repeats in the reference genome using RepeatModeler version 1.0.8 (Smit and Hubley 2015), which utilized RepeatMasker version open-4.0.7 (Smit *et al.* 2013) with the Repbase RepeatMasker libraries version 20170127 (Jurka 1998, 2000; Jurka *et al.* 2005; Bao *et al.* 2015), RMBlast version 2.2.28 (Smit *et al.* 2015), tandem repeats finder (TRF) version 4.09 (Benson 1999), RECON version 1.08 (Bao and Eddy 2002), and RepeatScout version 1.0.5 (Price *et al.* 2005). We first performed a homology-based masking of repeats and low-complexity regions and then implemented a second round of masking using the *de novo* repeat model we had generated using RepeatMasker version open-4.0.7 with the Repbase RepeatMasker libraries version 20170127, the repeat databases of the DFAM library version 2.0 (Hubley *et al.* 2016), TRF version 4.09, NCBI BLAST+ version 2.7.1 (Camacho *et al.* 2009), RMBlast version 2.2.28, and HMMER version 3.1b2 (Eddy 1998; http://hmmer.org). We created a browser extensible data (BED) formatted file of the repeat and low-complexity regions using seqtk version 1.2-r94 (Li 2016), sorted them using sort (GNU coreutils) version 8.25 (Haertel and Eggert 2016), and then used BEDTools version 2.25.0 (Quinlan and Hall 2010) to remove variants from our VCF file that fell in those regions. Finally, we removed variants at sites where the unfiltered read depth exceeded the mean plus five times the standard deviation, as suggested by the GATK documentation (https://software.broadinstitute.org/gatk/documentation/article.php?id=3225; Accessed 2018 Mar 16), using dp_cov_script.sh from SPOW-BDOW-introgression-scripts version 1.1.1 (Hanna *et al.* 2017b). We calculated the mean and standard deviation of the coverage depth across variant sites using DP_sample_calc.sh from genetics-tools version 1.0.0 (Hanna 2018b).

#### Population size and diversity statistics

After compressing and indexing our variant and repetitive region files using the bgzip and tabix tools from HTSlib version 1.7 (Li 2011; Davies *et al.* 2018), we treated each of the four samples as its own population and estimated the effective population size of each through time using SMC++ version 1.12.2.dev10+gaf60af6 (Terhorst *et al.* 2017) with the per-generation rate of collared flycatcher (*Ficedula albicollis*) (Smeds *et al.* 2016). We filtered the SMC++ files using GAWK version 4.2.0 and the GNU core utilities find version 8.25 (Decker *et al.* 2016), zcat version 8.25 (Gailly *et al.* 2016), and rm version 8.25 (Rubin *et al.* 2016). We converted our variant file format using BCFtools version 1.6 (Li *et al.* 2017) with HTSlib version 1.6 (Davies *et al.* 2017) and GAWK version 4.2.0. To estimate nucleotide diversity and population differentiation, we calculated π_Between_ (Nei and Li 1979) and the genome-wide fixation index (*F_ST_*) (Hudson *et al.* 1992) between *Strix occidentalis caurina* and *S. o. occidentalis* as well as between each of the population pairs using the tableFstPi script from diversity-divergence-stats version 1.0.0 (Wall and Hanna 2018) with cat (GNU coreutils) version 8.25 (Granlund and Stallman 2017). In addition, we calculated π_Within_ (Nei and Li 1979) for each of the subspecies and populations using the tablePiTheta script from diversity-divergence-stats version 1.0.0 with cat version 8.25. In order to check for evidence of recent inbreeding that could serve as an explanation for the elevated divergence of some population, we used GAWK version 4.2.0 and PLINK version 1.90b5.4 (Chang *et al.* 2015; Purcell and Chang 2018) to search for runs of homozygosity (ROH) in each individual. We only considered ROH that contained at least 100 SNPs, were at least 1 Mb long, and where the heterozygosity in the run was less than 10% of the genome-wide mean heterozygosity for the individual. Finally, we mapped our samples using QGIS version 2.18.2 (Quantum GIS Development Team 2017) with raster and vector files from Natural Earth (http://www.naturalearthdata.com) and a *S. occidentalis* range map from California Wildlife Habitat Relationships (Gould and CWHR Program 2008).

See full details of materials and methods at protocols.io (http://dx.doi.org/10.17504/protocols.io.qp2dvqe).

## Additional information and declarations

### Author contributions

Conceptualization: ZRH, JDW; Data Curation: ZRH, JPD, RCKB, JDW; Formal Analysis: ZRH, JDW; Funding Acquisition: JDW; Investigation: ZRH, JDW; Methodology: ZRH, JDW; Project Administration: ZRH, JDW; Resources: ZRH, JPD, RCKB, JDW; Software: ZRH, JDW; Supervision: JDW; Validation: ZRH; Visualization: ZRH, JDW; Writing – Original Draft Preparation: ZRH; Writing – Review & Editing: ZRH, JPD, RCKB, JDW

### Data availability

The raw sequences for *Strix occidentalis* samples CAS:ORN:99418, CDFWN13249, and CAFWN14089 are available from NCBI (SRA run accessions SRR6048848, SRR6048828, and SRR6048829, respectively).

## Acknowledgements

We thank J. Mark Higley and Aaron Pole of Hoopa Valley Indian Reservation Tribal Forestry; WildCare, San Rafael; Krysta Rogers, Wildlife Investigations Laboratory, California Department of Fish and Wildlife; and the California Academy of Sciences for graciously providing us with genetic samples.

## Supporting information

**Table S1. Variant site coverage values for each sample.** The site coverage is the average sequence coverage across all variant sites in our final variant set after filtering. “SD” stands for “standard deviation”.

**Table S2. Homozygous fraction of genome.** The total length of all runs of homozygosity (ROH) expressed as a fraction of the length of the reference genome for each population.

**Table S3. Specimen institution data.** We here provide information regarding the collections that archive specimens of the *Strix occidentalis* individuals we utilized in this study.

**Table S4. Genomic sequence details.** We here provide sample identifier information, including the NCBI Sequence Read Archive (SRA) run accessions of the raw genomic sequences we analyzed in our study.

## References

Bao, Z., and S. R. Eddy, 2002 Automated De Novo Identification of Repeat Sequence Families in Sequenced Genomes. Genome Res. 12: 1269–1276.

Bao, W., K. K. Kojima, and O. Kohany, 2015 Repbase Update, a database of repetitive elements in eukaryotic genomes. Mob. DNA 6: 1–6.

Barrowclough, G. F., and S. L. Coats, 1985 The demography and population genetics of owls, with special reference to the conservation of the spotted owl (*Strix occidentalis*), pp. 74–85 in Ecology and management of the spotted owl in the Pacific Northwest. Gen. Tech. Rep. PNW-185., U.S. Department of Agriculture, Forest Service, Pacific Northwest Forest and Range Experiment Station, Portland, OR.

Barrowclough, G. F., J. G. Groth, L. A. Mertz, and R. J. Gutiérrez, 2005 Genetic structure, introgression, and a narrow hybrid zone between northern and California spotted owls (*Strix occidentalis*). Mol. Ecol. 14: 1109–1120.

Barrowclough, G. F., R. J. Gutiérrez, and J. G. Groth, 1999 Phylogeography of Spotted Owl (*Strix occidentalis*) Populations Based on Mitochondrial DNA Sequences: Gene Flow, Genetic Structure, and a Novel Biogeographic Pattern. Evolution 53: 919–931.

Barrowclough, G. F., R. J. Gutiérrez, J. G. Groth, J. E. Lai, and D. F. Rock, 2011 The Hybrid Zone between Northern and California Spotted Owls in the Cascade—Sierran Suture Zone. The Condor 113: 581–589.

Benson, G., 1999 Tandem repeats finder: a program to analyze DNA sequences. Nucleic Acids Res. 27: 573–580.

Bolger, A. M., M. Lohse, and B. Usadel, 2014 Trimmomatic: a flexible trimmer for Illumina sequence data. Bioinformatics 30: 2114–2120.

Bryant, W. E., 1889 A catalogue of the birds of Lower California, Mexico, pp. 237–320 in Proceedings of the California Academy of Sciences, Second, California Academy of Sciences, San Francisco.

Camacho, C., G. Coulouris, V. Avagyan, N. Ma, J. Papadopoulos et al., 2009 BLAST+: architecture and applications. BMC Bioinformatics 10: 421.

Chang, C. C., C. C. Chow, L. C. Tellier, S. Vattikuti, S. M. Purcell et al, 2015 Second-generation PLINK: rising to the challenge of larger and richer datasets. GigaScience 4: 7.

Chi, T. Y., 2006 Genetic characterization of four populations in two subspecies of spotted owl [M.S.]: San Jose State University.

Clark, P. U., A. S. Dyke, J. D. Shakun, A. E. Carlson, J. Clark et al., 2009 The Last Glacial Maximum. Science 325: 710–714.

Davies, R., J. C. Randall, S. A. McCarthy, J. Bonfield, M. O. Pollard et al, 2017 HTSlib. Version 1.6. [Accessed 2018 Mar 20]. Available from: https://github.com/samtools/htslib.

Davies, R., J. C. Randall, S. A. McCarthy, J. Bonfield, M. O. Pollard et al, 2018 HTSlib. Version 1.7. [Accessed 2018 Mar 19]. Available from: https://github.com/samtools/htslib.

Davis, J. N., and G. I. Gould Jr., 2008 California spotted owl (*Strix occidentalis occidentalis*), pp. 227–233 in California Bird Species of Special Concern: A ranked assessment of species, subspecies, and distinct populations of birds of immediate conservation concern in California, edited by W. D. Shuford and T. Gardali. Studies of Western Birds 1, Western Field Ornithologists and California Department of Fish and Game (Sacramento, California), Camarillo, California.

Decker, E., D. MacKenzie, J. Plett, T. Wood, M. Rendell et al, 2016 find (GNU coreutils). Version 8.25. [Accessed 2018 Mar 19]. Available from: http://www.gnu.org/software/coreutils/coreutils.html.

DePristo, M. A., E. Banks, R. Poplin, K. V. Garimella, J. R. Maguire et al, 2011 A framework for variation discovery and genotyping using next-generation DNA sequencing data. Nat. Genet. 43: 491–498.

Dugger, K. M., E. D. Forsman, A. B. Franklin, R. J. Davis, G. C. White et al, 2015 The effects of habitat, climate, and Barred Owls on long-term demography of Northern Spotted Owls. The Condor 118: 57–116.

Eddy, S. R., 1998 Profile hidden Markov models. Bioinforma. Oxf. Engl. 14: 755–763.

Forsman, E. D., R. G. Anthony, J. A. Reid, P. J. Loschl, S. G. Sovern et al, 2002 Natal and Breeding Dispersal of Northern Spotted Owls. Wildl. Monogr. 1–35.

Free Software Foundation, 2017 GNU Awk.

Funk, W. C., E. D. Forsman, T. D. Mullins, and S. M. Haig, 2008 Introgression and dispersal among spotted owl (*Strix occidentalis*) subspecies. Evol. Appl. 1: 161–171.

Gailly, J., M. Adler, J. Meyering, and P. Eggert, 2016 Gzip (GNU coreutils).

Gould, G., and CWHR Program, 2008 Spotted Owl Range - CWHR B270 [ds897]. [Accessed 2018 May 10]. Available from: https://www.wildlife.ca.gov/Data/CWHR.

Granlund, T., and R. M. Stallman, 2017 cat (GNU coreutils).

Gutiérrez, R. J., 1994 Changes in the Distribution and Abundance of Spotted Owls During the Past Century. Stud. Avian Biol. 15: 293–300.

Gutiérrez, R. J., A. B. Franklin, and W. S. Lahaye, 1995 Spotted Owl (*Strix occidentalis*). The Birds of North America Online (A. Poole, Ed.). Ithaca: Cornell Lab of Ornithology. [Accessed 2016 Oct 1]. Retrieved from the Birds of North America Online: https://birdsna.org/Species-Account/bna/species/spoowl.

Gutiérrez, R. J., P. N. Manley, and P. A. Stine, 2017 The California spotted owl: current state of knowledge. Gen Tech Rep PSW-GTR-254 Albany CA US Dep. Agric. For. Serv. Pac. Southwest Res. Stn. 254:.

Haertel, M., and P. Eggert, 2016 sort (GNU coreutils).

Haig, S. M., T. D. Mullins, and E. D. Forsman, 2004 Subspecific relationships and genetic structure in the spotted owl. Conserv. Genet. 5: 683–705.

Haig, S. M., R. S. Wagner, E. D. Forsman, and T. D. Mullins, 2001 Geographic variation and genetic structure in Spotted Owls. Conserv. Genet. 2: 25–40.

Hanna, Z. R., 2018a Adapter sequences used for trimming of genomic sequences in the assembly of the Northern Spotted Owl (Strix occidentalis caurina) genome assembly version 1.0. Version 1.0.0. Zenodo.

Hanna, Z. R., 2018b genetics-tools. Version 1.0.0. Zenodo.

Hanna, Z. R., J. B. Henderson, A. B. Sellas, J. Fuchs, R. C. K. Bowie et al, 2017a Complete mitochondrial genome sequences of the northern spotted owl (*Strix occidentalis caurina*) and the barred owl (*Strix varia*; Aves: Strigiformes: Strigidae) confirm the presence of a duplicated control region. PeerJ 5: e3901.

Hanna, Z. R., J. B. Henderson, and J. D. Wall, 2017b SPOW-BDOW-introgression-scripts. Version 1.1.1. Zenodo.

Hanna, Z. R., J. B. Henderson, J. D. Wall, C. A. Emerling, J. Fuchs et al, 2017c Northern Spotted Owl (*Strix occidentalis caurina*) Genome: Divergence with the Barred Owl (*Strix varia*) and Characterization of Light-Associated Genes. Genome Biol. Evol. 9: 2522–2545.

Hanna, Z. R., J. B. Henderson, J. D. Wall, C. A. Emerling, J. Fuchs et al, 2017d Supplemental dataset for Northern Spotted Owl (Strix occidentalis caurina) genome assembly version 1.0. Zenodo.

Henke, A. L., 2005 Spotted owl (Strix occidentalis) microsatellite variation in California [M.S.]: San Jose State University.

Hubley, R., R. D. Finn, J. Clements, S. R. Eddy, T. A. Jones et al, 2016 The Dfam database of repetitive DNA families. Nucleic Acids Res. 44: D81–D89.

Hudson, R. R., M. Slatkin, and W. P. Maddison, 1992 Estimation of Levels of Gene Flow from DNA Sequence Data. Genetics 132: 583–589.

Jurka, J., 2000 Repbase Update: a database and an electronic journal of repetitive elements. Trends Genet. 16: 418–420.

Jurka, J., 1998 Repeats in genomic DNA: mining and meaning. Curr. Opin. Struct. Biol. 8: 333–337.

Jurka, J., V. V. Kapitonov, A. Pavlicek, P. Klonowski, O. Kohany et al, 2005 Repbase Update, a database of eukaryotic repetitive elements. Cytogenet. Genome Res. 110: 462–467.

Keane, J. J., 2017 Threats to the viability of California spotted owls, pp. 185–238 in The California Spotted Owl: Current State of Knowledge, edited by R. J. Gutiérrez, P. N. Manley, and P. A. Stine. General Technical Report PSW-GTR-254, U.S. Department of Agriculture, Forest Service, Pacific Southwest Research Station, Albany, California.

Li, H., 2013 Aligning sequence reads, clone sequences and assembly contigs with BWA-MEM. ArXiv:1303.3997 Q-Bio. [Accessed 2016 Feb 16].

Li, H., 2016 Seqtk. Version 1.2-r94. [Accessed 2018 Mar 16]. Available from: https://github.com/lh3/seqtk.

Li, H., 2011 Tabix: fast retrieval of sequence features from generic TAB-delimited files. Bioinformatics 27: 718–719.

Li, H., B. Handsaker, P. Danecek, S. McCarthy, J. Marshall et al, 2017 BCFtools. Version 1.6. [Accessed 2018 Mar 20]. Available from: https://github.com/samtools/bcftools.

McKenna, A., M. Hanna, E. Banks, A. Sivachenko, K. Cibulskis et al, 2010 The Genome Analysis Toolkit: A MapReduce framework for analyzing next-generation DNA sequencing data. Genome Res. 20: 1297–1303.

Miller, M. P., T. D. Mullins, E. D. Forsman, and S. M. Haig, 2017 Genetic differentiation and inferred dynamics of a hybrid zone between Northern Spotted Owls (*Strix occidentalis caurina*) and California Spotted Owls (*S. o. occidentalis*) in northern California. Ecol. Evol. 7: 6871–6883.

Montgomery, C. A., J. Brown Gardner M., and D. M. Adams, 1994 The Marginal Cost of Species Preservation: The Northern Spotted Owl. J. Environ. Econ. Manag. 26: 111–128.

Nei, M., and W. H. Li, 1979 Mathematical model for studying genetic variation in terms of restriction endonucleases. Proc. Natl. Acad. Sci. U. S. A. 76: 5269–5273.

Noon, B. R., and C. M. Biles, 1990 Mathematical Demography of Spotted Owls in the Pacific Northwest. J. Wildl. Manag. 54: 18–27.

Price, A. L., N. C. Jones, and P. A. Pevzner, 2005 De novo identification of repeat families in large genomes. Bioinformatics 21: i351–i358.

Purcell, S. M., and C. C. Chang, 2018 PLINK. Version 1.90b5.4. [Accessed 2018 Apr 12]. Available from: https://www.cog-genomics.org/plink/1.9.

Quantum GIS Development Team, 2017 Quantum GIS Geographic Information System. Open Source Geospatial Foundation Project. [Accessed 2017 Sep 16]. Available from: http://qgis.org.

Quinlan, A. R., and I. M. Hall, 2010 BEDTools: a flexible suite of utilities for comparing genomic features. Bioinformatics 26: 841–842.

Rubin, P., D. MacKenzie, R. M. Stallman, and J. Meyering, 2016 rm (GNU coreutils). Version 8.25. [Accessed 2018 Mar 19]. Available from: http://www.gnu.org/software/coreutils/coreutils.html.

Schoenherr, A. A., 1992 A Natural History of California. Berkeley: University of California Press.

Smeds, L., A. Qvarnström, and H. Ellegren, 2016 Direct estimate of the rate of germline mutation in a bird. Genome Res. 26: 1211–1218.

Smit, A. F. A., and R. Hubley, 2015 RepeatModeler Open-1.0. [Accessed 2018 Mar 16]. Available from: http://www.repeatmasker.org.

Smit, A., R. Hubley, and P. Green, 2013 RepeatMasker Open-4.0. [Accessed 2018 Mar 16]. Available from: http://www.repeatmasker.org.

Smit, A., R. Hubley, and National Center for Biotechnology Information, 2015 RMBlast. [Accessed 2016 Oct 1]. Available from: http://www.repeatmasker.org/RMBlast.html.

Tempel, D. J., R. J. Gutiérrez, and M. Z. Peery, 2017 Population distribution and trends of California spotted owls, pp. 75–108 in The California Spotted Owl: Current State of Knowledge, edited by R. J. Gutiérrez, P. N. Manley, and P. A. Stine. General Technical Report PSW-GTR-254, U.S. Department of Agriculture, Forest Service, Pacific Southwest Research Station, Albany, California.

Terhorst, J., J. A. Kamm, and Y. S. Song, 2017 Robust and scalable inference of population history from hundreds of unphased whole genomes. Nat. Genet. 49: 303–309.

USDA Forest Service, 1992 Final Environmental Impact Statement on Management for the Northern Spotted Owl in the National Forests. USDA Forest Service, National Forest System: Portland, Oregon. 2 vol.

Van der Auwera, G. A., M. O. Carneiro, C. Hartl, R. Poplin, G. del Angel et al, 2013 From FastQ data to high confidence variant calls: the Genome Analysis Toolkit best practices pipeline. Curr. Protoc. Bioinforma. 11: 11.10.1–11.10.33.

Wall, J. D., and Z. R. Hanna, 2018 diversity-divergence-stats. Version 1.0.0. Zenodo.

